# Differentiation of Acute Disseminated Encephalomyelitis from Multiple Sclerosis Using a Novel Brain Lesion Segmentation and Classification Pipeline

**DOI:** 10.1101/2024.07.23.604829

**Authors:** Osama Radi, Aiden Huang, Kira Murakami

**Affiliations:** Jeddah Knowledge School Jeddah, Saudi Arabia 23435; Acton-Boxborough Regional High School Massachusetts, United States 01720; Dulles High School Texas, United States 77478

## Abstract

Multiple Sclerosis (MS) is a chronic autoimmune disease affecting the central nervous system, while Acute Disseminated Encephalomyelitis (ADEM) is a sudden, often monophasic inflammatory condition of the brain and spinal cord. Only 17% of ADEM cases are correctly diagnosed on the first visit due to overlapping clinical and radiological presentations with Multiple Sclerosis (MS) [1]. Both ADEM and MS are demyelinating diseases, meaning they cause brain lesions by damaging the myelin sheath, leading to scar tissue that disrupts nerve signals [2]. Previous machine learning pipelines have differentiated Neuromyelitis Optica Spectrum Disorder (NMOSD) (a different demyelinating disease) from MS and ADEM from NMOSD based on MRI imagery with varying accuracies [3, 4]. Our novel Classifier for Demyelinating Disease (CDD) pipeline is the first to differentiate ADEM from MS using MRI imagery. It does this in two stages: a segmentation stage which creates segmentation masks of the lesions and a classification stage to classify them as either ADEM or MS. Additionally, we introduce a novel ADEM dataset from open-access medical case reports. The CDD pipeline achieves an accuracy of 90.0% on our validation dataset, making it a potentially viable diagnostic tool in the future. All data and code is available here. ^2^

## 1 Introduction

There is a critical need in the accurate differentiation between ADEM and MS due to the significant implications of mistreatment and misdiagnosis [5]. Radiologists often face challenges in distinguishing between these two conditions due to overlapping clinical and radiological features, leading to potential diagnostic inaccuracies [6]. Both MS and ADEM present with brain lesions of a similar type. A study conducted by Takahashi, Hayakawa and Abe showed that only 17% of ADEM cases were diagnosed correctly at the first visit [1].

Within the past 5 years, demyelinating disease classification using statistical machine learning has become a topic of interest in the field of computational neurology. In 2020, Wang et al. developed the first breakthrough in demyelinating disease classification. In this influential paper, a method of differentiating MS from NMOSD based on 3 dimensional Brain MRI images using convolutional neural networks was detailed. This model achieved an accuracy of 0.725 [3]. A second breakthrough was reached when Zhou et al. passed MRI images into a U-Net segmentation algorithm to separate the healthy matter from the lesions before putting the images through a CNN. In this study, they were able to differentiate NMOSD from ADEM with an accuracy of 95.55% [4]. Several approaches for lesion segmentation involve UNet [7], UNet++ [8], LinkNet [9], and 3D V-Net [10]. However, many segmentation methods struggle with the challenges of low contrast, fuzzy cell boundaries, and variable image quality involved with MRI scans of MS and ADEM, resulting in inaccurate diagnosis [11].

A pipeline differentiating ADEM from MS has yet to be developed; the CDD pipeline is the first of its kind in that manner. Additionally, the CDD pipeline is the first demyelinating disease classifying pipeline which utilizes the YOLOv8 segmentation algorithm in order to separate the brain lesions from the healthy brain matter [12].

The segmentation stage of the CDD pipeline is powered by the YOLOv8x segmentation model trained on ground truth brain lesion segmentation masks developed by three qualified neuroradiologists [13]. The classification stage is powered by a ResNet50 model pretrained on ImageNet then trained on the segmentation masks of MS and ADEM images created from the segmentation stage [14]. The MS data is gathered from the dataset present in the following paper [15]. Due to the disease’s rarity, there are no readily available datasets containing ADEM imagery. For that reason, the ADEM data is taken from 20 different case reports of patients with ADEM. This is done with explicit patient consent, as all patients consented to having their data publicly available. We contribute both a novel pipeline which can differentiate ADEM from MS with a high degree of accuracy, as well as a novel dataset containing T2-Flair MRI images of ADEM.

## 2 Methodology

**Figure 1:**
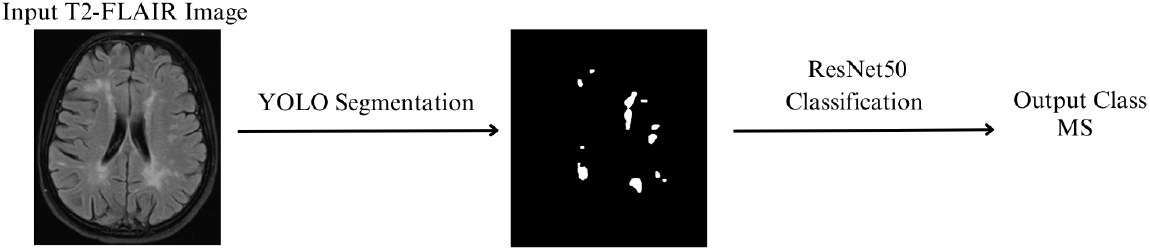
Diagram of the CDD pipeline outlining the segmentation and classification stages

### 2.1 Data Collection and Processing

The development of CDD involved meticulous data collection for both the segmentation and classification stages. For the segmentation phase, which uses a YOLOv8x model, T2-FLAIR brain MRI images of patients with brain lesions and ground truth segmentation masks were required. These masks were created by three highly qualified neuroradiologists to ensure precise annotations. The dataset for this phase was sourced from the Mendeley data repository [13]. For the classification phase, which employs a pretrained ResNet50 model through transfer learning, T2-FLAIR brain MRI images of patients diagnosed with either MS or ADEM were used. MS images were obtained from the dataset in the following paper [15]. Due to the rarity of ADEM, images were sourced from various different case reports, resulting in a varied and comprehensive dataset necessary for accurate classification. In total, 52 images were collected, with 38.5% used for validation data. The case reports which the ADEM data is compiled from are listed here. ^3^ We compiled the dataset of ADEM data from open-access case reports, which are publicly available and do not require additional patient consent for secondary use in research, in compliance with ethical guidelines and open-access policies.

Initially, the images and segmentation masks for training the YOLOv8 segmentation model were obtained in .nii format. Using the nibabel Python module, these images were converted to .jpg format to ensure compatibility with the segmentation model [33]. Additionally, the ground truth segmentation masks were formatted into the YOLO-compatible .txt format using the COCO2YOLO tool [34]. For the ResNet50 classification algorithm, uniform image size was essential. Thus, all T2-FLAIR brain MRI images for MS and ADEM cases were cropped to a standardized size of 390 × 442 pixels, ensuring consistency and effective learning in the classification model.

### 2.2 Segmentation

The segmentation phase of the project focused on training a YOLOv8x segmentation model to accurately identify and delineate brain lesions from T2-FLAIR brain MRI images [12]. The training process involved optimization over 300 epochs using a dataset comprised of training images and validation images, each paired with corresponding ground truth segmentation masks. These images and masks were sourced from the Mendeley dataset, specifically curated for T2-FLAIR brain MRI images of patients with brain lesions [13]. The YOLOv8 segmentation algorithm was selected due to its previous use in segmentation of tumors caused by breast cancer [35].

### 2.3 Classification

In the classification phase, a ResNet50 model pretrained on ImageNet underwent transfer learning using the segmentation masks generated by the YOLOv8 segmentation model [14, 12]. These masks, delineating lesions in T2-FLAIR brain MRI images, trained the ResNet50 model to classify the MRI images, distinguishing between MS and ADEM based on segmented regions. After training, the model’s performance was evaluated using a subset segmentation masks for validation, ensuring its ability to accurately classify MRI images across new data. The ResNet50 classication algorithm was selected due to its previous use in classifying pneumonia using segmented images [36].

## 3 Results

### 3.1 Segmentation

**Figure 2:**
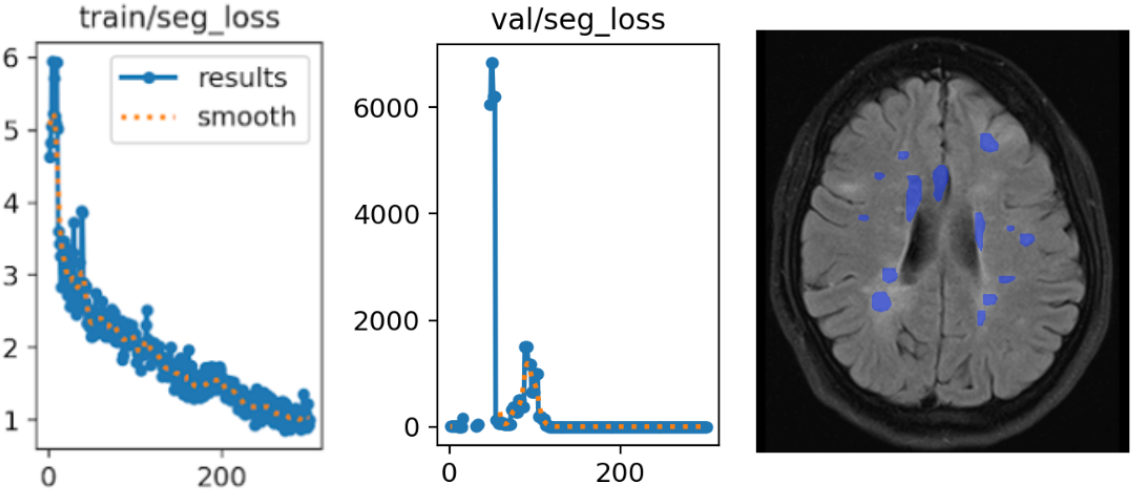
The segmentation loss plotted against number of epochs, the validation loss plotted against the segmentation loss, and a predicted segmentation mask from the validation data

The results of the segmentation phase indicate that the YOLOv8 model achieved a low segmentation loss of 1.0054 and a box loss of 0.668. These metrics suggest a high degree of accuracy in the model’s ability to identify and delineate brain lesions within the T2-FLAIR MRI images. The low segmentation loss reflects the model’s precision in matching the predicted segmentation masks to the ground truth masks, while the low box loss indicates effective localization of the lesions.

### 3.2 Classification

In the classification phase, the ResNet50 model demonstrated a notable accuracy of 0.9 in distinguishing between Acute Disseminated Encephalomyelitis (ADEM) and Multiple Sclerosis (MS). Despite the relatively small size of the training dataset, it was carefully balanced, containing an equal number of images for both ADEM and MS. Importantly, the testing images were entirely novel to the model, ensuring that none of these images were part of the training data. This high accuracy, achieved under stringent testing limitations, underscores the model’s effectiveness and potential utility in accurately classifying these neurological conditions based on T2-FLAIR brain MRI images.

## 4 Discussion

In our study, we were able to achieve an accuracy of 90.0% in differentiating between ADEM and MS with our new CDD framework. This paper introduces a novel method in differentiating these diseases.

The segmentation stage clearly shows how the pipeline is learning based off the criteria which are commonplace when classifying ADEM from MS. The Callen MS-ADEM critera states that MS shows multiple small, well-demarcated, periventricular lesions at various stages with ring/nodular enhancement, while ADEM presents larger, confluent, poorly demarcated, uniformly enhancing lesions in a single episode, often involving subcortical regions [37]. The clear presence of these features in the segmentation masks suggests that our pipeline is differentiating these images based on these general criteria.

The high accuracy rate achieved by the CDD pipeline indicates its potential ability to serve as a far more reliable method to classify ADEM than in current clinical practices, in which only 17% of ADEM cases were diagnosed accurately in the patient’s first visit [1].

While utilizing machine learning to differentiate ADEM from MS is a novel subject, papers have discussed the differentiation of other demyelinating diseases via machine learning in the past. In 2020, a study conducted by Wang et al. was able to achieve an accuracy of 72.5% in differentiating MS from NMOSD, another demyelinating disease, through the use of a 3d CNN architecture [3]. Our CDD framework builds off this research and is based on a recent 2024 study conducted by Zhou et al. which utilized a U-Net segmentation algorithm to identify and segment lesions and passed these segmented images into a CNN to be classified. In this study, Zhou et al. achieved an accuracy of 95.55% in differentiating NMOSD from ADEM [4]. Our CDD pipeline similarly utilizes a segmentation algorithm to mask the lesions followed by the use of a CNN to differentiate the diseases based on the segmented images. However, in our CDD framework, we introduced the novel use of the YOLOv8 model to segment brain lesions followed by the application of a pre-trained ResNet50 model to classify the diseases, resulting in highly accurate classification.

While we were able to achieve a high accuracy in differentiation between ADEM and MS, we believe that this pipeline can further be improved through the use of more data in training the model. This addition would prevent the model from generalizing the data. Additionally, the utilization of more recent YOLO models such as YOLOv9 for lesion segmentation could also improve accuracy. Finally, utilizing clinical presentation data such as fever or infection may further increase the accuracy of the model.

## 5 Conclusion

We contribute the novel CDD pipeline to differentiate Acute Disseminated Encephalomyelitis (ADEM) from Multiple Sclerosis (MS) as well as a novel ADEM dataset. The CDD pipeline segments lesions present in MRI scans through the use of the YOLOv8x segmentation model and classifies these segmented images as either ADEM or MS via a pretrained ResNet50 model through transfer learning. We conclude that the CDD pipeline achieves a high accuracy of 90.0% in differentiating between ADEM and MS. Given the distinct treatments for ADEM and MS, the ability to accurately diagnose these diseases is crucial. The need for accurate differentiation is further amplified by the common misclassification of the two diseases due to their similar and overlapping features [38]. The improper diagnosis of these diseases can lead to inappropriate treatment, resulting in worse patient outcomes [39]. The high classification accuracy achieved by the CDD framework substantially improves reliability in clinical diagnosis, which enhances patient recovery by allowing appropriate treatment. The use of a ResNet50 model allowed us to avoid retraining a CNN model from scratch and resulted in us achieving a high accuracy in spite of limited data availability.

## Footnotes

2 https://github.com/ozzux/CDD

3 [16, 17, 18, 19, 20, 21, 22, 23, 24, 25, 26, 27, 28, 29, 30, 31, 32]

## Notes

https://github.com/ozzux/CDD

https://www.kaggle.com/datasets/buraktaci/multiple-sclerosis

https://data.mendeley.com/datasets/8bctsm8jz7/1

### Competing Interest Statement

The authors have declared no competing interest.

